# Estimating inflation in GWAS summary statistics due to variance distortion from cryptic relatedness

**DOI:** 10.1101/164939

**Authors:** Dominic Holland, Chun-Chieh Fan, Oleksandr Frei, Alexey A. Shadrin, Olav B. Smeland, V. S. Sundar, Ole A. Andreassen, Anders M. Dale

## Abstract

Cryptic relatedness is inherently a feature of large genome-wide association studies (GWAS), and can give rise to considerable inflation in summary statistics for single nucleotide polymorphism (SNP) associations with phenotypes. It has proven difficult to disentangle these inflationary effects from true polygenic effects. Here we present results of a model that enables estimation of polygenicity, mean strength of association, and residual inflation in GWAS summary statistics. We show that there is substantial residual inflation in recent large GWAS of height and schizophrenia; correcting for this reduces the number of independent genome-wide significant loci from the reported values of 697 for height and 108 for schizophrenia to 368 and 61, respectively. In contrast, a larger GWAS of educational attainment shows no residual inflation. Additionally, we find that height has a relatively low polygenicity, with approximately 8k SNPs having causal association, more than an order of magnitude less than has been reported. The residual inflation in GWAS summary statistics can be corrected using the standard genomic control procedure with the estimated residual inflation factor.

## INTRODUCTION

Population structure in the context of Genome-wide association studies (GWAS) is usually modeled as a combination of two distinct types (Astle and Balding, 2009): population stratification, where the GWAS sample involves subsamples from different ethnicities or groups with different genetic ancestry; and (2) cryptic kinship or hidden family structure, where individuals assumed to be unrelated are in fact related. As GWAS involve increasingly large sample sizes, the effects of these sources of structure, especially in combination, become increasingly important (Price et al., 2010). In particular, if uncorrected they can profoundly affect GWAS summary statistics (z-scores for single nucleotide polymorphism (SNP) associations with a phenotype), though in distinct ways: stratification is a source of confounding bias, leading to shifts in the estimates of the true effect sizes of SNPs, while cryptic relatedness will lead to variance distortion of the z-scores. For highly polygenic traits, it can be difficult to disentangle causal effects – their number and size – from apparent effects arising from population structure (Yang et al., 2011). Pure stratification can be modeled and corrected using ancestry principal components as covariates in the association analysis (Price et al., 2006), and is routinely performed in GWAS. The effects of pure variance distortion of z-scores for phenotypes with very low polygenicity can largely be corrected by the standard genomic control procedure – rescaling the z-scores so that the median z^2^ is that of the centered χ^2^ distribution with 1 degree of freedom (Devlin and Roeder, 1999). Mixed regression models (or linear mixed models) relying on raw genotype data have also been developed to account for both cryptic kinship and population stratification (Kang et al., 2010, 2008; Zhou and Stephens, 2012; Loh et al., 2015), though there are advantages and pitfalls in their application (Yang et al., 2014).

In the published literature, the inflation of GWAS summary statistics is generally assumed to be negligible (Schizophrenia Working Group of the Psychiatric Genomics Consortium, 2014; Wood et al., 2014). Yet despite significant effort to correct for population structure in the asociation analyses, and the development of methods to detect residual inflation in published summary statistics (Bulik-Sullivan et al., 2015), we report here on evidence from different lines of enquiry of significant residual inflation in the summary statistics from recent large GWAS of height and schizophrenia. In contrast, summary statistics from a larger GWAS for educational attainment show no sign of residual inflation. We provide estimates for the inflation, which can be corrected using the standard genomic control procedure (Devlin and Roeder, 1999).

## METHODS

### Causal Model for GWAS Summary Statistics

We recently proposed a model probability distribution function (PDF) for the distribution of GWAS z-scores, with three parameters: the polygenicity, π_1_; the variance of the true causal effect sizes 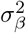 (the variance of the true association β coefficients); and the residual variance of the z-scores, 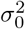, which in the absence of population structure is unity (Holland et al., 2017). This model takes into account the detailed linkage disequilibrium (LD) structure of the SNPs. 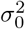 can be thought of as the true genomic control factor: it quantifies inflation in z-scores from factors other than causal polygenicity. The traditional genomic control factor, λ, gives the inflation above the null hypothesis value of z-scores from all sources, including causal polygenicity which causes it to increase with sample size.

## Data

As in our previous work (Holland et al., 2017), we calculated SNP minor allele frequency (MAF) and LD between SNPs using the 1000 Genomes phase 3 data set for 503 samples of European ancestry (Consortium et al., 2015, 2012; Sveinbjornsson et al., 2016), for a total of n_*snp*_=11,015,833 SNPs. We analyzed GWAS summary statistics for participants with European ancestry for: height (Wood et al., 2014) with median SNP sample size 251,783 (min 50,003; max 253,280) for a total of 2,036,013 SNPs; schizophrenia (Schizophrenia Working Group of the Psychiatric Genomics Consortium, 2014), with 35,476 cases and 46,839 controls (N_*ef f*_ 4/(1/N_*cases*_ +1/N_*controls*_) = 76, 326), with imputation of SNPs using the 1000 Genomes Project reference panel (1000 Genomes Project Consortium, 2010) for a total of 5,369,285 genotyped and imputed SNPs; and educational attainment (Okbay et al., 2016), measured as the number of years of schooling completed, with 328,917 samples and a total of 5,361,110 SNPs, available at https://www.thessgac.org In performing the model fit as described in Holland et al. (2017), we pruned synonomnous SNPs, i.e., those with LD r^2^ > 0.8.

## RESULTS

Figure 1 shows QQ plots for the z-scores for height, schizophrenia, and educational attainment, along with estimates based on the model PDF. In all cases, the model fit (yellow) closely tracks the data (dark blue). For height we find that the residual variance distortion of the z-scores, the “true” genomic control factor, is 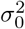= 1.50, while for schizophrenia it is 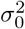= 1.18. Although educational attainment has a much larger sample size (∼ 329k), and is the most polygenic of the three phenotypes (π_1_ = 7.7 × 10^-3^ ), its GWAS summary statistics show no sign of inflation due to variance distortion: 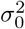= 1.01. In all cases, however, the model estimate, λ̂, of the traditional genomic control factor almost exactly predicts the value λ (given in the figure) calculated directly from the pruned data: λ̂ =1.79, 1.38, and 1.17, for height, schizophrenia, and education, respectively. (Note that genomic control values for pruned data are always lower than for unpruned data. E.g., the overall traditional genomic control factor for the entire set of available z-scores for height is λ = 1.92.) The high value of 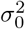 for height results in the visually steep rise of the QQ plot near the origin.

**Figure 1:**
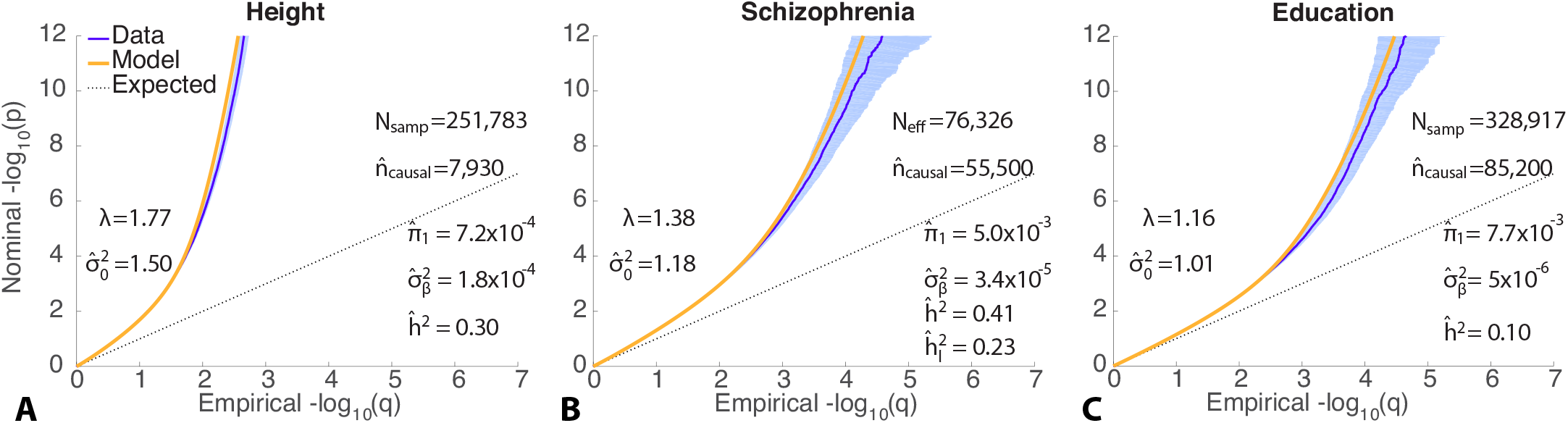
QQ plots of z-scores for (A) height, (B) schizophrenia, and (C) educational attainment, (dark blue, 95% confidence interval in light blue) with model prediction (yellow). *q* is the proportion of z-scores exceeding a p-value threshold given by *p*. The dashed line is the expected QQ plot under null (no SNPs associated with the phenotype). *λ* is the overall traditional genomic control factor for the pruned data (which is accurately predicted by the model: λ̂ = 1.79, 1.38, and 1.17, for (A), (B), and (C), respectively). The three estimated model parameters are: polygenicity, *π*̂_1_; discoverability, 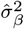; and SNP association *χ*^2^-statistic inflation factor, 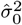. *h*̂ ^2^ is the estimated narrow-sense *β* 0 chip heritability (reexpressed as 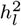 on the liability scale for schizophrenia assuming a prevalence of 1% in adult populations), and *n*̂_*causal*_ is the estimated number of causal SNPs. *n_snp_* = 11, 015, 833 is the total number of SNPs, whose LD and MAF underlie the model; the GWAS z-scores are for subsets of these SNPs. *N_samp_* is the sample size, expressed as *N_ef f_* for schizophrenia – see text. Reading the plots: on the vertical axis, choose a p-value threshold (more extreme values are further from the origin), then the horizontal axis gives the proportion of SNPs exceeding that threshold (higher proportions are closer to the origin). Figure 5 in Supporting Material shows the full y-axis, up to *p* = 10^−150^ for height.

We also estimated residual inflation in three other ways. Figure 2 shows a QQ plot for height restricted to SNPs with low heterozygosity (H=2p(1-p), where p is either allele frequency) and low total LD (TLD, the sum of LD r^2^ with neighboring SNPs thresholded at 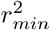 = 0.05, where r is the minor allele count correlation). These are SNPs which are least likely to show large z^2^ values from polygenic association; the traditional genomic control value for these SNPs is expected to reflect sources other than causal polygenicity, i.e., 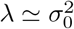. Restricting to SNPs with 0.01 < H ≤ 0.15 and 1 ≤ TLD ≤ 40, we find that λ = 1.43. We also carried out linear regression on the mean z^2^ for SNPs in equally-spaced total LD bins (for TLD ≤ 150), and for SNPs in equally-spaced heterozygosity bins. Since SNP z-scores can be decomposed into a component, arising from SNP–phenotypic association and an environmental component, *z* = √ NH β_eff_ + ∈ where N is the sample size and β_*ef f*_ is the association effect arising through LD with neighboring causal SNPs, and 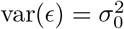(Holland et al., 2017), in the limit of low TLD and/or low H it follows that var 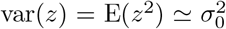. Thus, for both TLD regression and H regression of z^2^, the intercepts should approximately equal 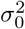. The intercepts were 1.56 and 1.53, respectively. In the absence of non-polygenic sources of inflation, these values would be expected to be close to 1.

**Figure 2.**
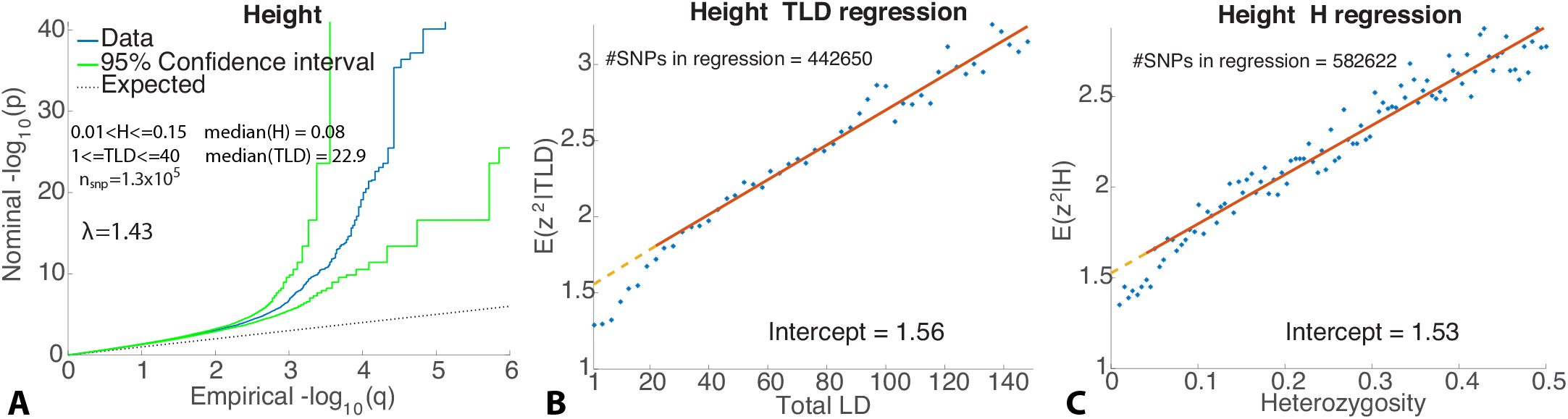
(A) QQ plot for height (in blue, with 95% confidence interval in green) restricted to SNPs with low total LD (TLD) and heterozygosity (H; *n_SN P_* is the number of such SNPs), giving *λ* = 1.43 (approximately equal to 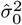 in Fig. 1(A)), and in reasonable agreement with the intercept from regressing *z*^2^ on (B) total LD, and (C) heterozygosity. Thus, different approaches to estimating residual inflation converge on approximately the same value, indicating the presence of population structure that has not been accounted for in the published summary statistics.

For schizophrenia, a similar analysis gave λ = 1.17 for the low H and low TLD SNPs, Figure 3(A), in agreement with 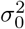in Figure 1(B), while the intercepts from regressing z^2^ on TLD and H were both 1.16, Figure 3(C).

**Figure 3.**
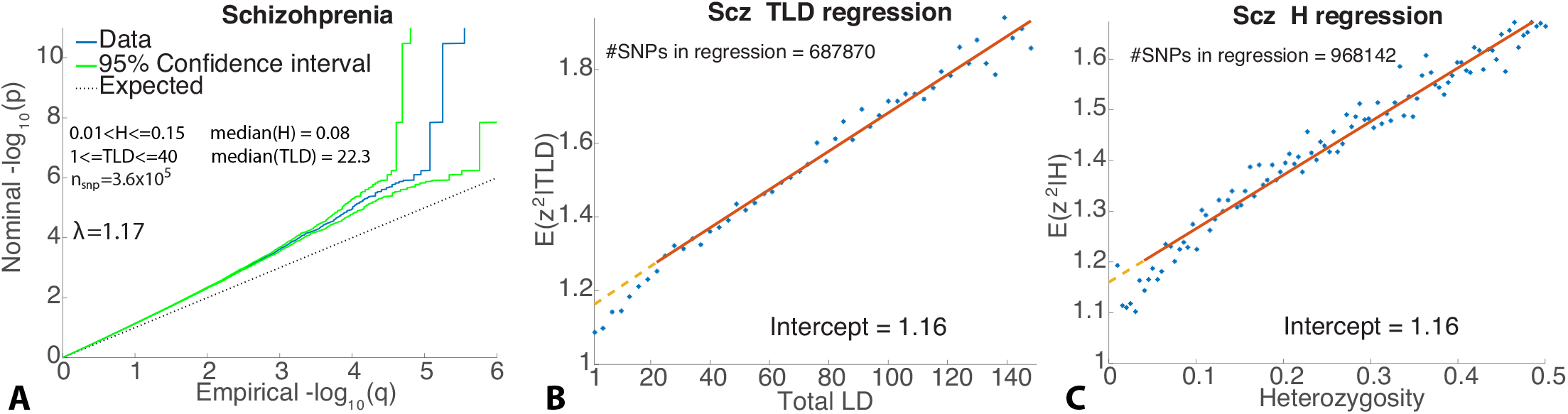
(A) QQ plot for schizophrenia (in blue, with 95% confidence interval in green) restricted to SNPs with low total LD (TLD) and 2 heterozygosity (H; *n_SN P_* is the number of such SNPs), giving *λ* = 1.17 (approximately equal to 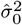 in Fig. 1(B)), and in agreement with the intercept from regressing *z*^2^ on (B) total LD, and (C) heterozygosity for schizophrenia.

Taking 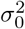 = 1.5 to indicate the true residual inflation in the height z-scores (i.e., assuming the z^2^-values are too large by 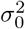 = 1.5), correcting for this is equivalent to requiring the given p-values to exceed (be less than) *pt* = 2.4×10^−11^ in order to pass genome-wide significance – more stringent than *p_GWAS_* = 5 × 10^−8^, which would apply if there were no inflation. Then, instead of the 697 SNPs reaching genome-wide significance (Supplementary Table 1 in Wood et al. (2014)), there would be only 368. For schizophrenia, taking 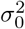 = 1.18 to indicate the true residual inflation, the given p-values would need to be less than *p_t_* = 3.2×10^−9^ to reach significance, resulting in only 61 instead of the 108 independent loci reaching genome-wide significance (Supplementary Table 3 in Schizophrenia Working Group of the Psychiatric Genomics Consortium (2014)).

For height we find that the polygenicity is π_1_ = 7.2 × 10^−4^, almost an order of magnitude less than that of schizophrenia, while the variance of the effect sizes is 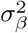 = 1.8 × 10^−4^, almost an order of magnitude greater than the value for schizophrenia. Constraining σ_0_ = 1, the model gives a much poorer fit (indicated by the large mismatch between λ = 1.77 and λ̂ = 1.60), with elevated π_1_ = 3.3 × 10^-3^ (Supplementary Figure 7).

A power analysis (see (Holland et al., 2017)) showed that for height 46% of the narrow-sense heritability arising from common SNPs is explained by genome-wide significant SNPs at the current sample size (N=251,783).

## DISCUSSION

Despite great effort – and much success – in genome-wide association studies to provide SNP summary statistics that are free of inflation arising from population structure, we find evidence that there remains considerable inflation in the summary statistics for two well-studied phenotypes: height and schizophrenia. It is not clear what exactly is the source of this inflation, though cryptic relatedness is likely the cause, nor how it survived the procedures employed to mitigate it. Notably, a much larger GWAS for the more highly polygenic phenotype of educational attainment showed no signs of having inflated summary statistics.

For height, we find the true genomic control factor to be 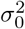 = 1.5. The inflation component of this (0.5) is more than half of the inflation component (0.92) arising from the overall traditional genomic control factor for the entire set of available z-scores, λ = 1.92. For schizophrenia, we find the true genomic control factor to be 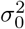 = 1.18, while the overall traditional genomic control factor for the entire set of available z-scores is λ = 1.55. In the GWAS reports for both cases (Wood et al., 2014; Schizophrenia Working Group of the Psychiatric Genomics Consortium, 2014), the inflation has been assumed to be negligible (i.e., with 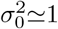).

If the height GWAS does indeed involve large inflation as indicated by 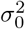 = 1.5, this conceivably could lead to a substantial over-estimate of the polygenicity of height, since many SNPs would appear to show association signal (even if below the threshold of genome-wide significance). A recent estimate puts the polygenicity at 3.8%, indicating that on the order of 400k SNPs have causal effects on height (Boyle et al., 2017); notably, the methods employed for this estimation did not allow for test statistic inflation (Stephens, 2016). Taking into account the apparent residual inflation indicated by 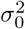, we find the polygenicity to be fifty times lower: 0.072%, for approximately only 8k causally-associated SNPs, suggesting that height is not in fact “omnigenic” (Boyle et al., 2017). Additionally, inflation in test statistics inflates the number of loci reaching genome wide significance. Thus, for height, of the 697 independent SNP identified in Wood et al. (2014) as causally associated with height, only 368 would actually reach genome-wide significance; for schizophrenia only 61 of the 108 independent loci identified in Schizophrenia Working Group of the Psychiatric Genomics Consortium (2014) would reach genome-wide significance.

In simulations we recently carried out using realistic LD structure (Holland et al., 2017), 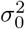 is close to 1.0 (indicating no variance distortion, and hence no inflation due to cryptic relatedness, as expected in HapGen (Su et al., 2011)), though there is a trend toward larger values for higher heritability and polygenicity. Thus, the very high inflation found for height and schizophrenia is unlikely to be an artifact of the causal model.

Our estimates of inflation in height and schizophrenia from our causal effects model are corroborated by intercept estimates from regressing z^2^ on total LD, and regressing z^2^ on heterozygosity, both of which give intercepts that are close to the respective 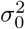 estimates, and additionally by calculating the traditional genomic control λ from SNPs selected for low total LD and low heterozygosity. In contrast, the LD Score regression intercept estimate for schizophrenia is only 1.07 (Bulik-Sullivan et al., 2015).

(Note that *H* and total LD are highly correlated – see Supplementary Fig. 6.) As *H* → 0, E(*z*^2^) = var(*z*) = 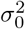. Thus, the intercept (at *H* = 0) from regressing z^2^ on *H* gives an estimate of 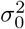. The intercept (at T LD = 1) from regressing z^2^ on T LD gives an estimate of the variance of z-scores for those SNPs uninfluenced by the effects of polygenicity through LD, and thus similarly gives an estimate of the residual inflation arising through bias in the data.

**Figure 4:**
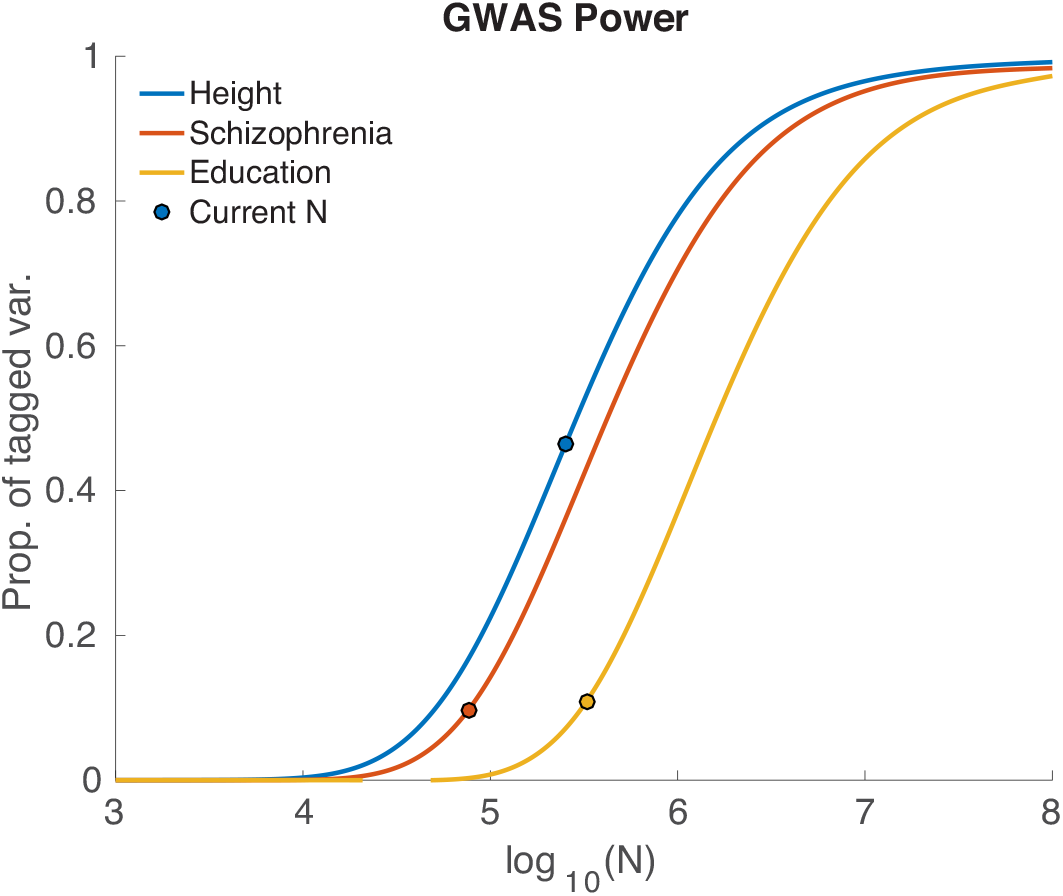
Proportion of narrow-sense chip heritability captured by genome-wide significant SNPs as a function of sample size, *N*. Left-to-right plot order is determined by decreasing 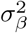. For current sample sizes, the proportions are: height, 0.46; schizophrenia, 0.096; educational attainment, 0.109.

**Figure 5:**
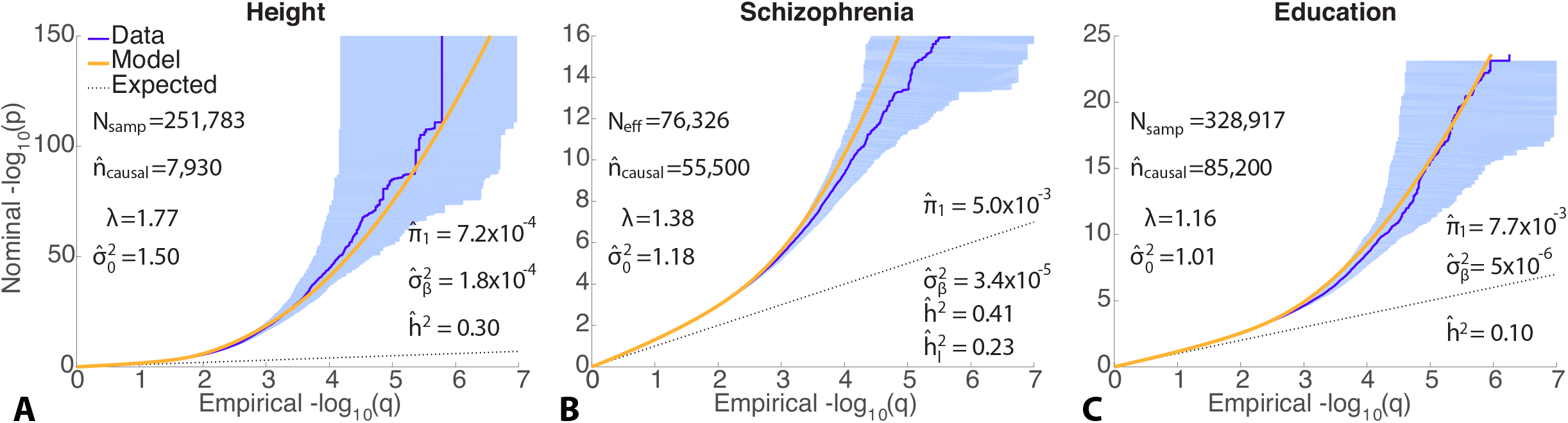
QQ plots of z-scores for (A) height, (B) schizophrenia, and (C) educational attainment. Similar to Figure 1 in the main paper, but with extended y-axis.

Our liability-scale heritability estimate for schizophrenia is substantially lower than the estimate from LD Score regression on the same overall PGC2 data set used here, 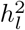 = 0.555 (Bulik-Sullivan et al., 2015). When the per-allele effect size, β, is independent of the MAF, LD Score regression will lead to a biased rotation of the regression line, increasing the slope, which is proportional to the estimated heritability, and decreasing the intercept, which gives the residual inflation. Thus, the discrepancy with LD Score regression in principle might arise due to the assumption in LD Score regression that effect sizes for variants are drawn independently from normal distributions with variance inversely proportional to heterozygosity (so that the variance explained per SNP is uncorrelated with T LD, since T LD is positively correlated with MAF). However, we show that the expected z^2^ increases linearly with H (as well as T LD), indicating that E(*β*^2^) = 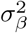is, to a first approximation, independent of MAF. Thus, the assumption of larger effects (β) for less common variants appears unwarranted, and would lead to both a lower estimate of residual inflation from population structure/cryptic relatedness and a higher estimate of heritability.

For height, the GWAS estimate of narrow sense heritability, calculated using GCTA applied to a subset of the raw genotype data, has been reported to be 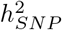 = 0.5 and that 32% of this (i.e., 16% of total phenotypic variation) is explained by genome-wide significant SNPs (Wood et al., 2014). Our estimate based on summary statistics for the same GWAS is 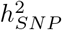 = 0.3, and that 46% of this (i.e., 14% of total phenotypic variation) is explained by genome-wide significant SNPs. Our heritability estimate uses the average heterozygosity over all SNPs (Holland et al., 2017). Conceivably, with the relatively small number of causal SNPs involved in height, this could be an underestimate for the causal SNPs themselves.

## CONCLUSION

In the new era of very large scale GWAS, it will be increasingly important to differentiate between the effects of polygenicity and inflation. Larger sample sizes will lead to larger genomic control factors, with the contributions from population structure becoming more pronounced as cryptic relatedness and stratification become larger features of the GWAS samples. We have shown here how these effects can be estimated from summary statistics. Where inflation exists, phenotypes that may have appeared to be highly polygenic or even omnigenic will be less so, and false discoveries become less likely.

**Figure 6.**
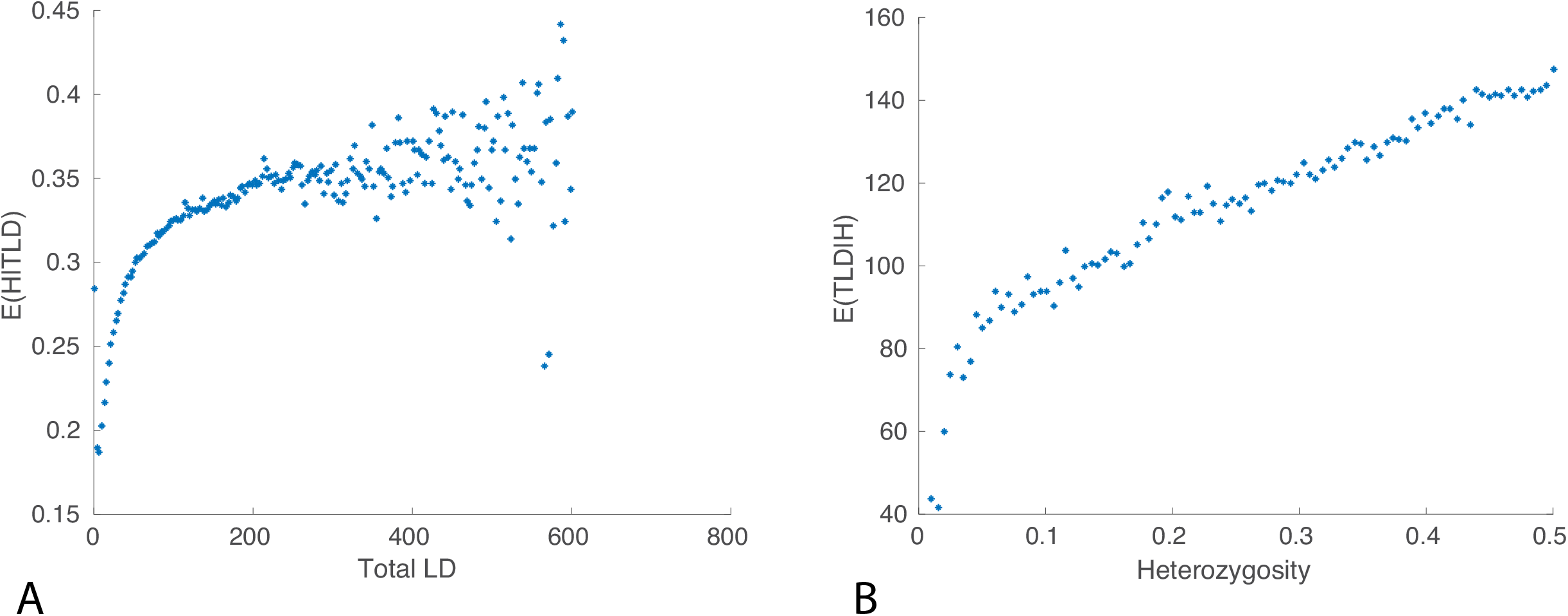
(A) Mean value of heterozygosity for given total LD (SNPs were binned based on TLD and the mean TLD for each bin plotted on the x-axis; the corresponding mean heterozygosity for SNPs in each bin was then plotted on the y-axis). (B) Mean value of total LD for given heterozygosity. Plots made for SNPs in the PGC2 schizophrenia GWAS; TLD and H calculated from 1000 Genomes phase 3 reference panel.

**Figure 7.**
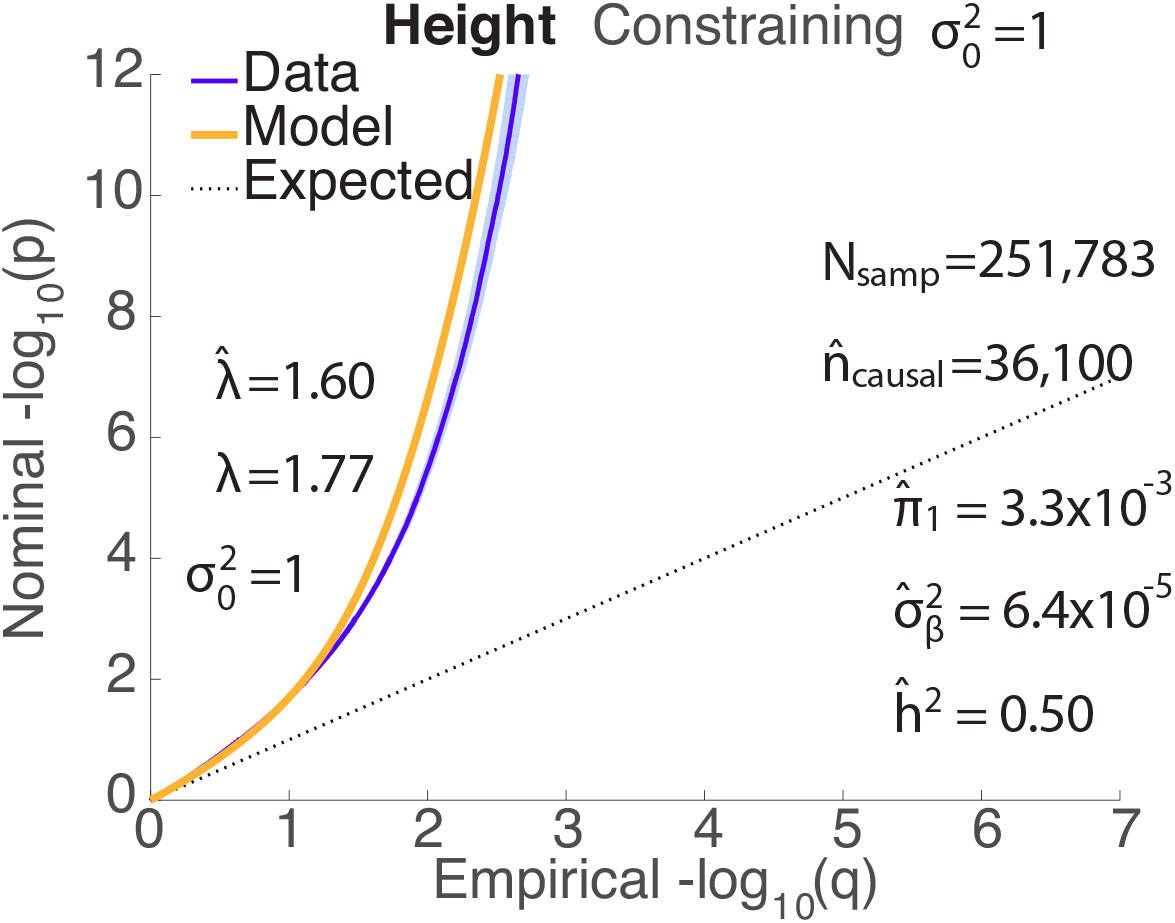
QQ plots of z-scores for height (dark blue, 95% confidence interval in light blue) with model prediction under the constraint *σ*_0_ = 1 (yellow). The dashed line is the expected QQ plot under null (no SNPs associated with the phenotype). *λ* is the overall traditional genomic control factor for the pruned data; with *σ*_0_ sonstrained to be 1, the model prediction of this quantity is *λ*̂ = 1.60. The two estimated model parameters are: polygenicity, *π*̂_1_; and discoverability, 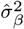. *h*̂ ^2^ is the estimated narrow-sense chip heritability, and *n*̂_*causal*_ is the estimated number of causal SNPs. *n_snp_* = 11, 015, 833 is the total number of SNPs, whose LD and MAF underlie the model; the GWAS z-scores are for subsets of these SNPs. *N_samp_* is the sample size. Compare Figure 1(A).

## Acknowledgments

We thank the Schizophrenia Working Group of the Psychiatric Genomics Consortium (PGC) for making available their GWAS summary statistics for schizophrenia, and the Social Science Genetic Association Consortium (SSGAC) for GWAS summary statistics on educational attainment.

## References

1000 Genomes Project Consortium, 2010. A map of human genome variation from population-scale sequencing. Nature 467 (7319), 1061–1073.

Astle, W., Balding, D. J., 2009. Population structure and cryptic relatedness in genetic association studies. Statistical Science, 451–471.

Boyle, E. A., Li, Y. I., Pritchard, J. K., 2017. An expanded view of complex traits: From polygenic to omnigenic. Cell 169 (7), 1177–1186.

Bulik-Sullivan, B. K., Loh, P.-R., Finucane, H. K., Ripke, S., Yang, J., Patterson, N., Daly, M. J., Price, A. L., Neale, B. M., of the Psychiatric Genomics Consortium, S. W. G., et al., 2015. Ld score regression distinguishes confounding from polygenicity in genome-wide association studies. Nature genetics 47 (3), 291–295.

Consortium,. G. P., et al., 2012. An integrated map of genetic variation from 1,092 human genomes. Nature 491 (7422), 56–65.

Consortium,. G. P., et al., 2015. A global reference for human genetic variation. Nature 526 (7571), 68–74.

Devlin, B., Roeder, K., Dec 1999. Genomic control for association studies. Biometrics 55 (4), 997–1004.

Holland, D., Fan, C.-C., Frei, O., Shadrin, A. A., Smeland, O. B., Sundar, V., Andreassen, O. A., Dale, A. M., et al., 2017. Estimating phenotypic polygenicity and causal effect size variance from GWAS summary statistics while accounting for inflation due to cryptic relatedness. bioRxiv, 133132.

Kang, H. M., Sul, J. H., Zaitlen, N. A., Kong, S.-y., Freimer, N. B., Sabatti, C., Eskin, E., et al., 2010. Variance component model to account for sample structure in genome-wide association studies. Nature genetics 42 (4), 348–354.

Kang, H. M., Zaitlen, N. A., Wade, C. M., Kirby, A., Heckerman, D., Daly, M. J., Eskin, E., 2008. Efficient control of population structure in model organism association mapping. Genetics 178 (3), 1709–1723.

Loh, P.-R., Tucker, G., Bulik-Sullivan, B. K., Vilhjalmsson, B. J., Finucane, H. K., Salem, R. M., Chasman, D. I., Ridker, P. M., Neale, B. M., Berger, B., et al., 2015. Efficient bayesian mixed-model analysis increases association power in large cohorts. Nature genetics 47 (3), 284–290.

Okbay, A., Beauchamp, J. P., Fontana, M. A., Lee, J. J., Pers, T. H., Rietveld, C. A., Turley, P., Chen, G.-B., Emilsson, V., Meddens, S. F. W., et al., 2016. Genome-wide association study identifies 74 loci associated with educational attainment. Nature 533 (7604), 539–542.

Price, A. L., Patterson, N. J., Plenge, R. M., Weinblatt, M. E., Shadick, N. A., Reich, D., 2006. Principal components analysis corrects for stratification in genome-wide association studies. Nature genetics 38 (8), 904–909.

Price, A. L., Zaitlen, N. A., Reich, D., Patterson, N., 2010. New approaches to population stratification in genome-wide association studies. Nature Reviews Genetics 11 (7), 459–463.

Schizophrenia Working Group of the Psychiatric Genomics Consortium, Jul 2014. Biological insights from 108 schizophrenia-associated genetic loci. Nature 511 (7510), 421–427.

Stephens, M., 2016. False discovery rates: a new deal. Biostatistics 18 (2), 275–294.

Su, Z., Marchini, J., Donnelly, P., 2011. Hapgen2: simulation of multiple disease snps. Bioinformatics 27 (16), 2304–2305.

Sveinbjornsson, G., Albrechtsen, A., Zink, F., Gudjonsson, S. A., Oddson, A., Ma´sson, G., Holm, H., Kong, A., Thorsteinsdottir, U., Sulem, P., et al., 2016. Weighting sequence variants based on their annotation increases power of whole-genome association studies. Nature genetics.

Wood, A. R., Esko, T., Yang, J., Vedantam, S., Pers, T. H., Gustafsson, S., Chu, A. Y., Estrada, K., Luan, J., Kutalik, Z., et al., 2014. Defining the role of common variation in the genomic and biological architecture of adult human height. Nature genetics 46 (11), 1173–1186.

Yang, J., Weedon, M. N., Purcell, S., Lettre, G., Estrada, K., Willer, C. J., Smith, A. V., Ingelsson, E., O’connell, J. R., Mangino, M., et al., 2011. Genomic inflation factors under polygenic inheritance. European Journal of Human Genetics 19 (7), 807–812.

Yang, J., Zaitlen, N. A., Goddard, M. E., Visscher, P. M., Price, A. L., 2014. Advantages and pitfalls in the application of mixed-model association methods. Nature genetics 46 (2), 100–106.

Zhou, X., Stephens, M., 2012. Genome-wide efficient mixed-model analysis for association studies. Nature genetics 44 (7), 821–824.

